# Emergent critical oscillations in motor cortex of Parkinson’s patients

**DOI:** 10.64898/2026.01.09.698590

**Authors:** Cheng Ly, J. Sam Sooter, Andrea K. Barreiro, Antonio J. Fontenele, Woodrow L. Shew

## Abstract

The dynamical state of cortical neural activity constrains the complexity of functions it can perform. A marginally stable dynamical state - called criticality - is thought to be beneficial for brain functions that require multiple time scales, broad dynamic range, and large information storage and transmission. A growing body of evidence suggests that criticality is a feature of healthy brain dynamics, but breaks down in certain brain disorders. Here we ask whether Parkinson’s disease incurs deviation from criticality compared to healthy controls. We analyze human resting state EEG activity in primary motor cortex. Parkinson’s patients exhibit prominent oscillatory brain activity in multiple frequency bands (low delta and high theta) that is not present in controls. Surprisingly, we find that these emergent oscillations are close to criticality, i.e., amplitude fluctuations with approximate temporal scale invariance. We compare traditional signatures of criticality and more principled measurements of proximity to criticality using our new approach based on information theory using temporal renormalization group. Our new approach and traditional methods agree, demonstrating that critical dynamics are not always associated with healthy states; Parkinson’s disease is associated with the emergence of near-critical oscillations in motor cortex.

**New and Noteworthy:** Brain function is thought to be optimal near critical states that border order and chaos. We investigate whether Parkinson’s disease disrupts this balance like in other diseases and pathologies. In resting state EEG of motor cortex, we show that people with Parkinson’s have rhythmic activity near the critical state. We show that critical dynamics can accompany disease, suggesting that being closer to criticality is not always a sign of healthy brain function.

## Introduction

What are the properties of cortical neural activity that confer its ability to perform healthy functions? One long-standing hypothesis posits that a healthy brain operates in a dynamical state near *criticality* - a special, marginally stable state imbued with a wide range of scale-invariant time scales and optimal computation (Shew and Plenz, 2013; Hengen and Shew, 2025). Indeed, evidence for criticality is associated with improved cognitive performance in humans (Müller et al., 2025; Xin et al., 2025; Ezaki et al., 2020) and multiple beneficial computational properties including efficient coding (Safavi et al., 2024), large dynamic range (Kinouchi and Copelli, 2006; Shew et al., 2009; Gautam et al., 2015), discrimination of sensory input (Clawson et al., 2017; Gautam et al., 2015), and more. If these properties of criticality are needed for healthy brain function, then unhealthy dysfunction may be associated with deviation from criticality. This has indeed been reported in multiple studies (Zimmern, 2020). For instance, Alzheimer’s disease (Montez et al., 2009; Ghassemkhani et al., 2025; McGregor et al., 2024), schizophrenia (Nikulin et al., 2012; Moran et al., 2019), depression (Linkenkaer-Hansen et al., 2005), and epilepsy (Fuscà et al., 2023) are associated with deviation from criticality compared to controls. However, the notion that healthy brain function requires closeness to criticality is challenged by some studies. For instance, sustained, focused attention seem to cause deviation from criticality (Irrmischer et al., 2018; Fagerholm et al., 2015). Here our primary goal was to determine how Parkinson’s disease impacts criticality in motor cortex. Motor cortex is a crucial area for voluntary movement and muscle control, functions that are severely impaired in Parkinson’s disease. We analyze a publicly available EEG dataset (Jackson et al., 2019; Swann et al., 2015; George et al., 2013) and ask whether motor cortex dynamics are closer to criticality for healthy control subjects or Parkinson’s patients.

To rigorously measure proximity to criticality, we use our newly-developed approach based on information theory and Gaussian autoregressive processes that relies on developing temporal Renormalization Group (**tRG**) theory to find all of the critical points (Sooter et al., 2026). This approach measures distance from criticality (*d*_2_) from time series data based on the nature of temporal fluctuations. We first show in a ground-truth model that critical dynamics can occur with prominent oscillations, i.e., ‘critical oscillations’. We then apply the tRG framework to EEG data for the first time, to our knowledge, and compare it to traditional methods of quantifying timescales from time series (Zeraati et al., 2024) including the decay times of autocorrelation function (**ACF**) (Müller and Meisel, 2023; Müller et al., 2025), and long-range temporal correlation via detrended fluctuation analysis (**DFA**) (Peng et al., 1994; Hardstone et al., 2012). However, we emphasize that our new approach rests on a mathematically rigorous definition of proximity to criticality, which is lacking in traditional methods (Tian et al., 2022).

We find that motor cortex resting state EEG activity in Parkinson’s patients is marked by the emergence of near critical oscillations in the delta and theta bands; oscillations in these bands not present in healthy controls. Two recent studies are consistent with our results, although we measure distance to criticality directly. Calvo et al. (2024) found in whole-brain human MEG data that control subjects were further from a chaotic point in more frequency bands than Parkinson’s, and Lee et al. (2024) found in whole-brain human EEG that control subjects had shorter timescales (measured by DFA) than Parkinson’s in the theta band in some brain regions. Our results indicate that critical dynamics are not always associated with healthy states; Parkinson’s disease results in critical oscillations.

## Results

### How oscillations can be critical

A common notion about oscillatory time-series activity is that it cannot have critical dynamics because of the presence of a characteristic timescale (the period of oscillation). Because one feature of physical systems at criticality is the *absence* of a characteristic timescale, this would seem to be a contradiction.

An alternative question to ask is whether the amplitude fluctuations of the oscillatory activity will display signatures of critical activity. This is referred to as *critical oscillations* (Palva and Palva, 2018). Neural activity within frequency bands is known to be relevant for neural computation (Buzśaki, 2006). In fact, using band-pass filtering and envelope extraction are general practice for large-spatial-scale time series data acquired through modalities such as EEG (Cohen, 2014) (see Fig 2A).

We show here that critical oscillations arise in a ground-truth model, and that the envelope fluctuations display the rich statistical properties associated with time series from critical systems. Because we have an analytically specified model, we can tune it closer to or further away from criticality; we also show that our new rigorous measure of proximity to criticality that we call d_2_ (see **Methods: Temporal Renormalization Group Theory** and results below for more details) successfully identifies the proximity to criticality and is better than other common methods.

Consider a linear rate model, with population activity vector 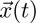 governed by:

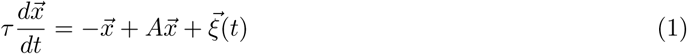

where *x_j_*(*t*) is the activity of the *j*^th^ neuron at time *t*, *A* is an *N × N* coupling (or adjacency) matrix with weights drawn from a Gaussian distribution with zero mean and variance of 2. Self-connections were removed by setting all diagonal elements *A_ii_* to zero. No additional structural constraints such as symmetry or Dale’s law were imposed. 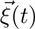 is uncorrelated white noise: 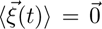, *⟨ξ_j_*(*t*)*ξ_k_*(*t^′^*)*⟩* = *τσ*^2^*δ_j,k_δ*(*t − t^′^*); we set *N* = 500 and *τ* = 50 ms; we down-sampled the simulated activity to 46 ms time bins so it is comparable to data (Pachitariu et al., 2026). This model has been widely used and insightful for studying chaos (Sompolinsky et al., 1988) and critical dynamics, for example by Hu and Sompolinsky (2022); Dahmen et al. (2019) who derived formulas for the spectrum of the covariance of activity (and studied dynamics near criticality with applications to experimental data). Also, Morales et al. (2023) used this model framework to infer the spectrum paired with phenomenological renormalization group applied to experimental data to compare scale-invariant statistics from recordings across many different brain regions, and most recently by Fontenele et al. (2026) to study hidden critical modes.

The spectrum of *A* determines whether this model exhibits critical dynamics, i.e., max *Re*(*λ*) = 1 corresponds to criticality, where *λ* are the eigenvalues of *A*. More generally, if *Re*(*λ_i_*) = 1, then the *i*^th^ mode is at criticality, and furthermore if *Im*(*λ_i_*) ≠ 0 then the *i*^th^ mode will display critical oscillations (Fontenele et al., 2026). Below we demonstrate: i) how critical oscillations can emerge, ii) our distance to criticality measure *d*_2_ (in bits/sec to quantify accumulation of evidence for ruling out being at criticality) applied to the entire population averaged activity in the model 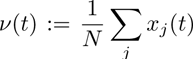, i.e, PSTH, is an excellent approximation to the ground truth assessment of proximity to criticality, iii) other methods (autocorrelation function (**ACF**) timescale, detrended fluctuation analysis (**DFA**); see Fig 1C and **Methods**) behave as expected at criticality even with prominent oscillations.

**Fig. 1.**
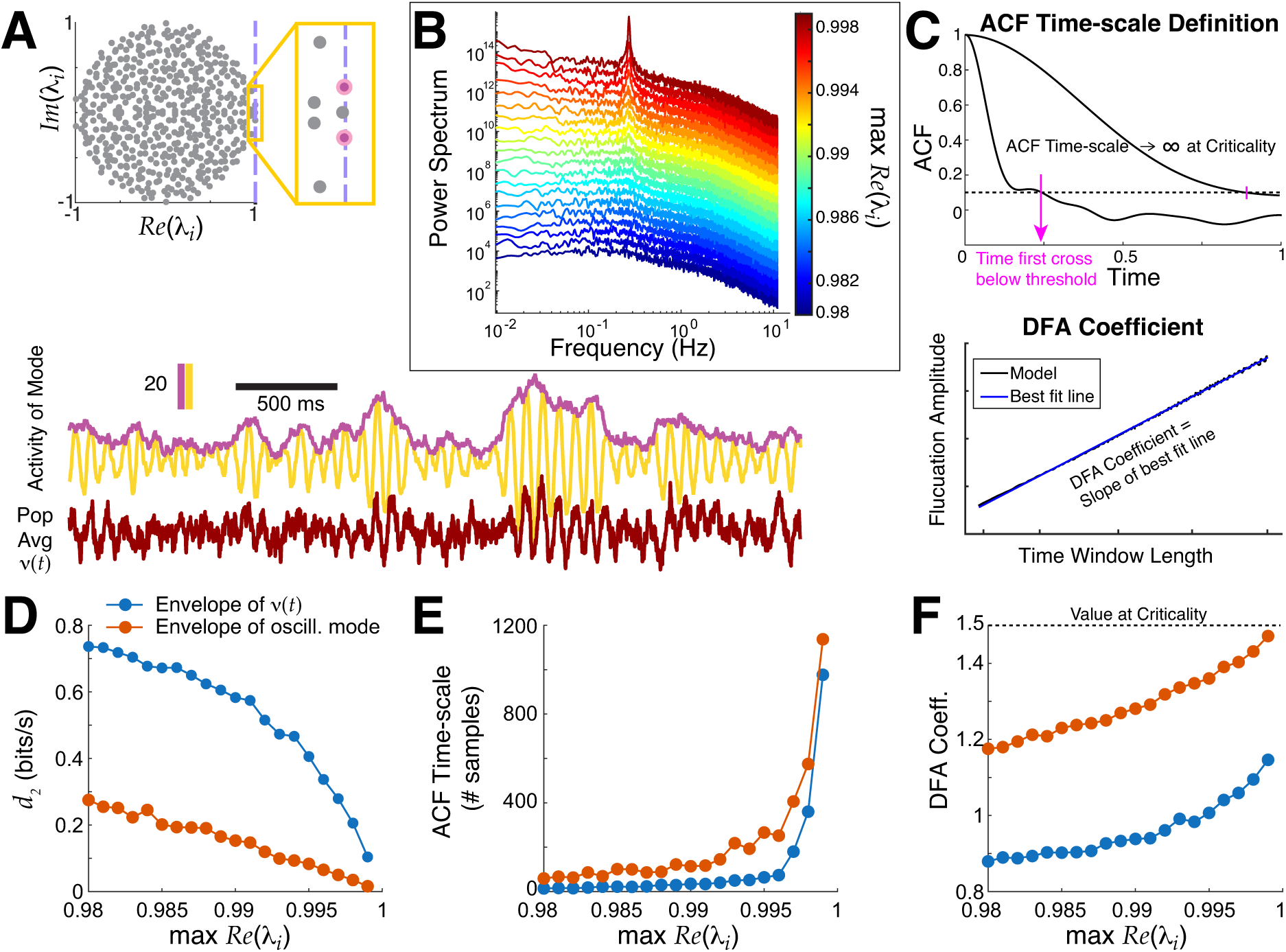
Critical oscillations in a ground truth model are well-captured by our methods. **A)** Top: spectrum of coupling matrix *A* near critical oscillations. Bottom: simple population average *ν*(*t*) (dark red bottom trace) does not have critical dynamics but the oscillatory mode (yellow) has critical dynamics in the amplitude fluctuations, revealed by taking the envelope (purple) (with max *Re*(*λ*) = 0.998). **B**) The power spectrum of *ν*(*t*) as the network approaches criticality has emergent oscillations (blue to red). **C**) Two traditional methods for measuring time-series data: ACF timescale (top) and DFA (bottom, no numbers/units for demo). A consequence of approaching critical dynamics is ACF timescale increases and DFA coefficient approaches 1.5. **D, E, F**) Our distance to criticality method *d*_2_, ACF timescale (number of samples; *ν*(*t*) was down-sampled to 46 ms time bins (Pachitariu et al., 2026)), and DFA coefficient calculations applied to *ν*(*t*) (blue) <u>all</u> well-approximate the underlying critical oscillations (blood orange) that are obscured in the entire population activity.

Our model study of proximity to criticality involves scaling all entries of the coupling matrix *A* systematically so that max *Re*(*λ*) *→* 1*^−^* (Fig 1A), which leads to rich network activity statistics (Fontenele et al., 2026). The power spectrum of the entire population averaged activity *ν*(*t*), i.e, PSTH, has a prominent peak as the network approaches criticality (Fig 1B), i.e., critical oscillations. This behavior is easily constructed by insuring the leading eigenvalue (max *Re*(*λ*)) is complex and has non-zero imaginary part; the power spectrum has a prominent peak because of the leading oscillatory mode even though there are contributions from other non-oscillatory modes. We apply the *d*_2_ method to the population firing rate *ν*(*t*) after band-pass filtering and envelope extraction, akin to what we will do in the EEG data. Figure 1D–F shows that these methods well-approximate the ground truth distance to criticality in the model. A consequence of (near) critical dynamics is that traditional measures for assessing lengths of timescales should diverge, which is precisely what we observe in the model with ACF timescale (the first time the ACF crosses below a threshold, Fig 1E); also, the DFA exponent increases (Fig 1F). Finally, the network activity *ν*(*t*) also exhibits avalanche statistics when very close to criticality (Fig 2 in (Fontenele et al., 2026)).

**Fig. 2.**
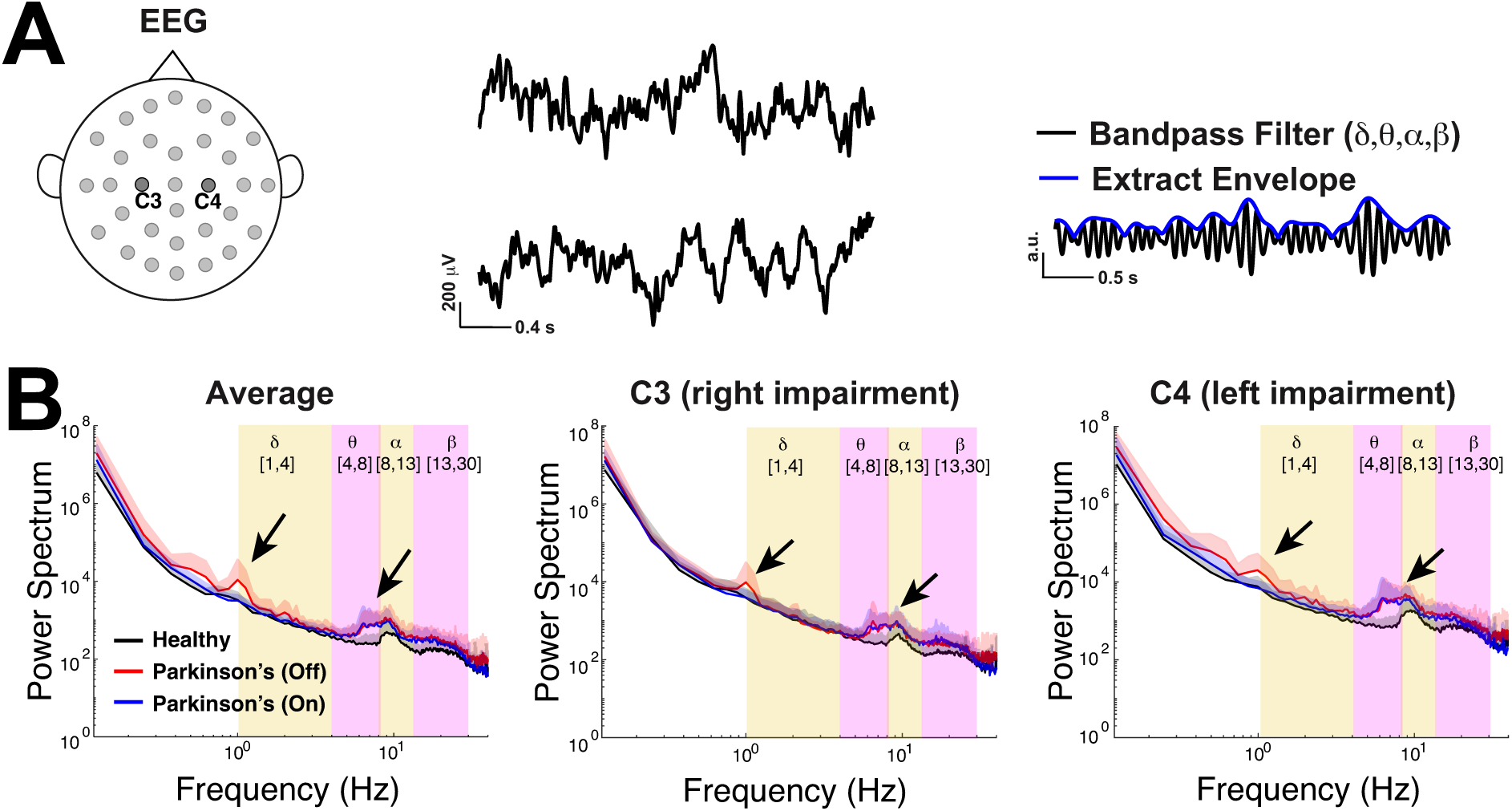
Emergent *δ* and *θ* oscillations in Parkinson’s patients. A) EEG time-series from two electrodes at the primary motor cortex on the left (C3) and right side (C4); right panel illustrates extracting the envelope of the band-passed EEG recording. **B**) The population-averaged power spectrum of the envelope of the EEG (without any band-pass filtering) for respectively the average of C3 and C4, as well as C3 and C4 individually, all exhibit peaks in the lower delta-band for Parkinson’s patients off medication (red), peaks in upper theta- to lower alpha-bands for Parkinson’s patients (on and off drugs), and a peak in the alpha-band for control (black). The shaded region above the curve corresponds to one standard deviation across the subjects.

We also observe that in this model, taking the whole population average obscures the presence of critical oscillations in the leading mode (blue curves are not the same as the blood orange curves in Fig 1D–F), but nevertheless these methods applied to the whole population monotonically track the ground truth (orange).

Taken together and considering that *d*_2_ is a rigorous and direct measure for distance to criticality, we conclude that using *d*_2_ is a better measure for quantifying critical oscillations in real data given that the ground truth is inaccessible. Having validated the use of distance to criticality in a model where the ground truth is known, we next applied the method to an EEG dataset.

### Application to EEG Parkinson’s data

We use freely available resting state EEG data (Fig 2A) collected by a lab in UCSD (Jackson et al., 2019; Swann et al., 2015; George et al., 2013) using a standardized format (Pernet et al., 2019; Appelhoff et al., 2019). Following previous studies (Jackson et al., 2019; Swann et al., 2015), we analyze data from the two electrodes positioned over left and right primary motor cortex (**M1**) labeled C3 and C4 (Fig 2A), an important brain region for motor planning and voluntary movement, functions that are impaired in these Parkinson’s patients. The dataset consists of 3 minute recording sessions from 16 healthy control subjects (**control**) and 15 subjects with Parkinson’s disease in two states: off drugs and on drugs to manage their symptoms. The Parkinson’s patients exhibited slight variability in severity of the disease as measured by Unified Parkinson’s Disease Rating Scale (**UPDRS**) III, but otherwise were not cognitively impaired compared to control subjects via Mini-Mental Status Exam (**MMSE**) or the North American Reading Test (**NAART**) (George et al., 2013). We use a common approach: applying a band-pass filter to the EEG data and subsequently extracted the amplitude envelope (Cohen, 2014) (Fig 2A right panel) to be used as the signal for all the analyses here (except Fig 2B). By studying fluctuations of the amplitude envelope, we can assess whether the oscillations at particular frequency bands are near or far from criticality. By definition, a critical oscillation will have amplitude fluctuations that are temporally scale invariant (Fontenele et al., 2026; Palva and Palva, 2018). This approach follows the long tradition of studying critical oscillations, more commonly assessed via long range temporal correlations (**LRTC**) (Linkenkaer-Hansen et al., 2001, 2005; Jackson et al., 2019; Hohlefeld et al., 2012, 2015) equivalent to DFA, an entity that is known to increase with critical dynamics.

This EEG data has power in select frequency bands (Fig 2B), a common observation in other resting state EEG data (Newson and Thiagarajan, 2019). Importantly, the timescales and dynamics in the broadband signal (Fig 2B) are distinct from the corresponding entities in band-passed power-envelope signal that is the main focus of our study. Figure 2B shows the population average (across 16 control subjects and 15 Parkinson’s patients) power spectrum of the envelope of the EEG data (y-axes on a log-scale) without band-pass filtering. Whether considering the average of both C3 and C4 (left panel), or a single electrode alone (C3 in middle, C4 in right panel), it is evident that control subjects on average (black curve) have peaks in their power spectrum in the alpha-band (8 to 13 Hz), a signature of healthy (Britton et al., 2016). Parkinson’s patients, however, have emergent oscillations in the lower *δ*-band (1 to 4 Hz) for patients off medication (red) and in the *θ*-band for both Parkinson’s subjects on (blue) and off medication.

We start by analyzing the timescales of fluctuations in oscillation amplitudes using two traditional tools: autocorrelation (see Fig 3A and Eq (2)), detrended fluctuation analysis (see Fig 4A and **Methods: Detrended Fluctuation Analysis**). Then, we compare to our new information theoretic method for measuring distance to criticality *d*_2_, (where tRG theory was used to find all of the scale-invariant fixed points) that goes beyond simple measures of timescales. For the control subjects, we simply use the average of both electrodes, but for Parkinson’s patients we use the electrode that corresponds to the side that subjects are reported to have physical limitations, see Table 3. The results reported in the main text (Figs 3–5) also hold when we use both electrodes in Parkinson’s patients, see Supplementary Text S1 and Figures S1, S2. Figure 3A shows the average autocorrelation function (over the number of subjects) in the four frequency bands (the alpha- and theta-bands zoomed-in to the right of the main axes) where the control subjects (black curves) have consistently faster decay than Parkinson’s patients (blue and red, x-axis at specific time lags is a log-scale). As in Fig 1C,E, we calculate the ‘ACF timescale’ by finding the first time a subject’s ACF falls below a chosen threshold (Fig 3B). The population ACF timescales exhibit statistically significant differences between healthy control and Parkinson’s patients, specifically the control subjects generally have <u>faster timescales</u> than Parkinson’s patients (both on and off drugs) in the delta- and theta-frequency bands (population summarized with box plots, Fig 3C), statistical significance was assessed with the Wilcoxon rank-sum test under the null hypothesis that the values were generated from the same probability distribution (see **Methods: Wilcoxon Rank-Sum Test** for details). When the differences were significant, the effect sizes are medium to large (see Table 1). There are no statistical differences between Parkinson’s patients on drugs versus off drugs (see Table 1; the population means are included in Fig 3D for completeness). Although commonly used, the autocorrelation function simply measures statistical correlation at a specific time lag averaged over the entire time series, in contrast to other measures that account for fluctuation trends as window sizes vary (DFA). Note that there are other methods for extracting timescales from the ACF, such as fitting an exponential function (Siegle et al., 2021; Li and Wang, 2022; Zeraati et al., 2022, 2024) or its variants (Zeraati et al., 2022; van Meegen and van Albada, 2021); but in this data, neither the population averages nor individual ACFs are well-fit by exponential functions.

**Fig. 3.**
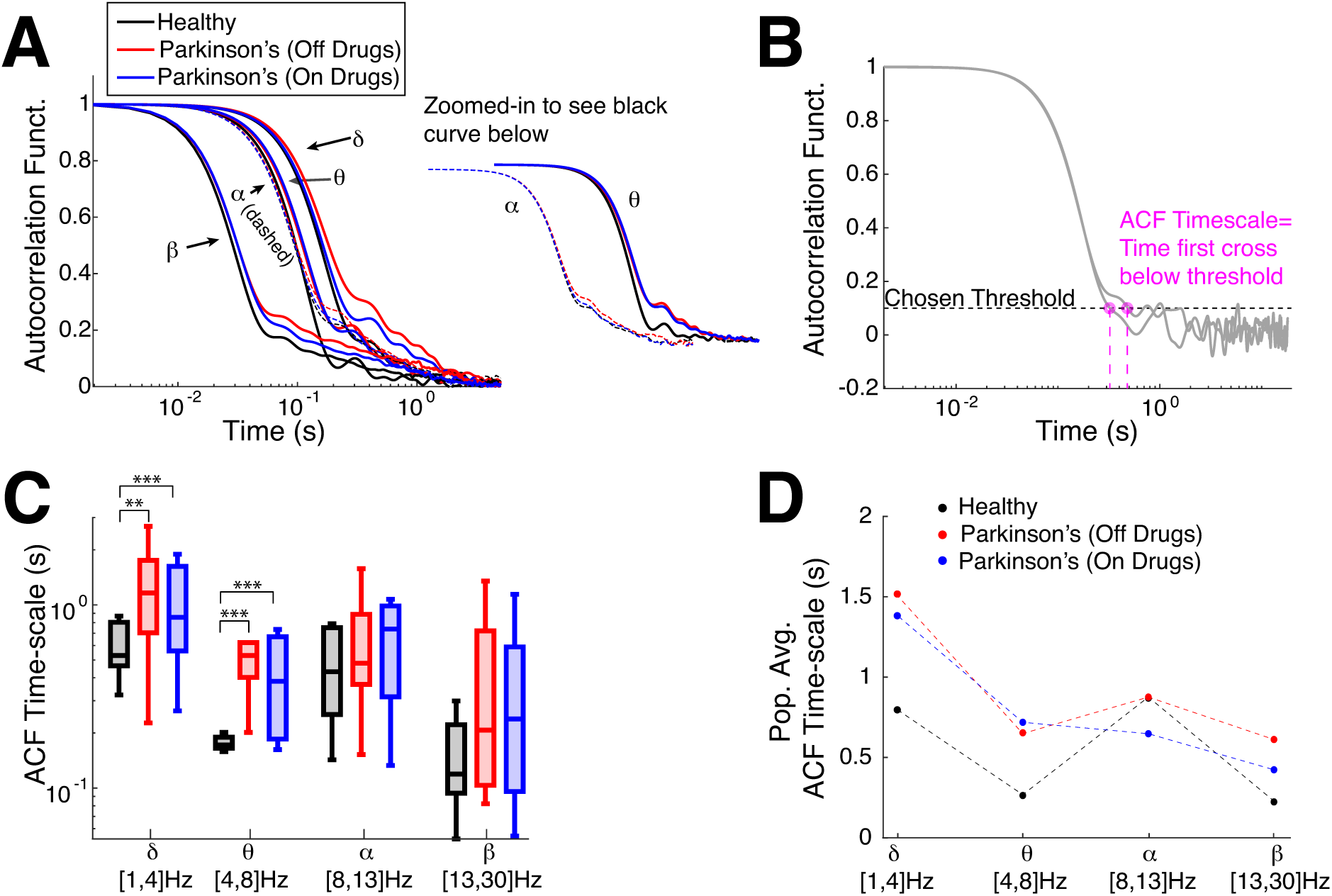
Emergent Parkinson’s oscillations have large autocorrelation time. **A)** The population-averaged ACF in Parkinson’s patients has longer timescales (red, blue: slower decay) in motor cortex EEG activity across all 4 frequency bands than healthy (control). ACF in alpha- and theta-band are zoomed-in and shifted for clarity. **B)** Example calculation of ACF timescale, i.e., a measure of ACF decay time, for 2 subjects; the first time where a subject’s ACF falls below a <u>chosen threshold of 0.1</u>. **C)** Summary of ACF timescale with box plots in different frequency bands shows that control subjects on average have faster ACF time decay (smaller timescale) than Parkinson’s patients for the lower frequency bands (delta and theta). The horizontal lines in the boxes represent inter-quartiles: 25^th^ percentile, median, and 75^th^ percentile. Difference in distributions are statistically significant measured by Wilcoxon rank-sum test (see Table 1 for details). **D**) The population means of ACF timescale are plotted for completeness.

**Fig. 4.**
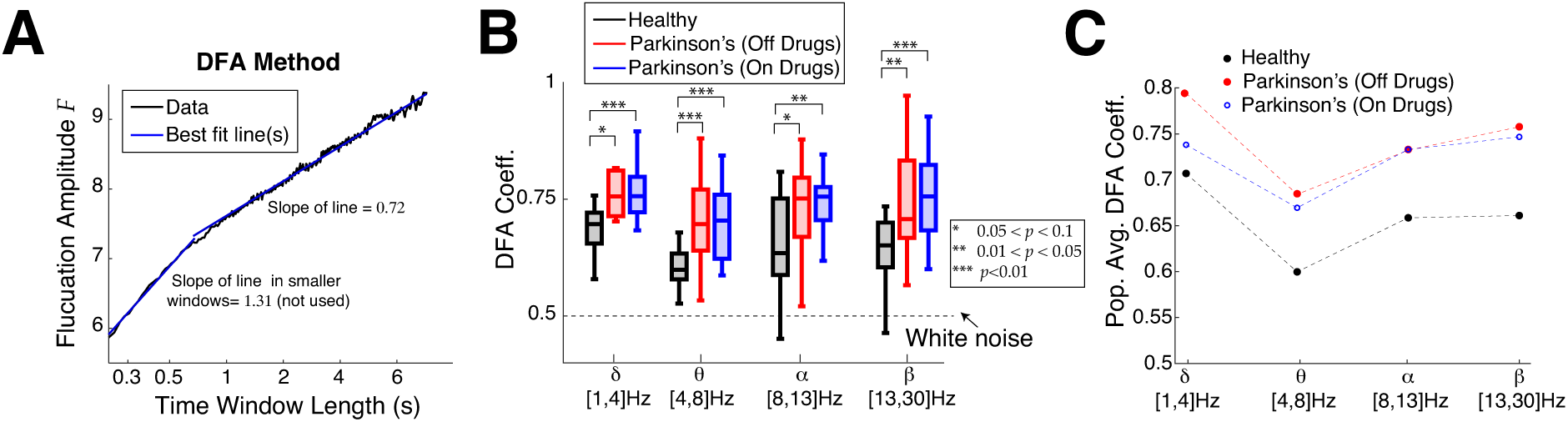
Emergent Parkinson’s oscillations have larger DFA exponents. **A)** Example DFA coefficient calculation (control subject 1 in delta-band) well-fit with 2 line segments, where <u>a choice for the time window</u> to use has to be made. When 2 lines are used, the slope of the right line segment for larger time windows is reported. **B)** Summary of DFA coefficients with box plots in different frequency bands is largely consistent with the ACF results (Fig 3C,D). Box plot convention are the same as in Figure 3C. The results are not as strong in the alpha-band. Difference in distributions are statistically significant measured by Wilcoxon rank-sum test (see Table 2 for details). **C**) The population means of DFA coefficients are plotted to clearly illustrate that control subjects are further from criticality/scale-invariance than Parkinson’s patients.

**Fig. 5.**
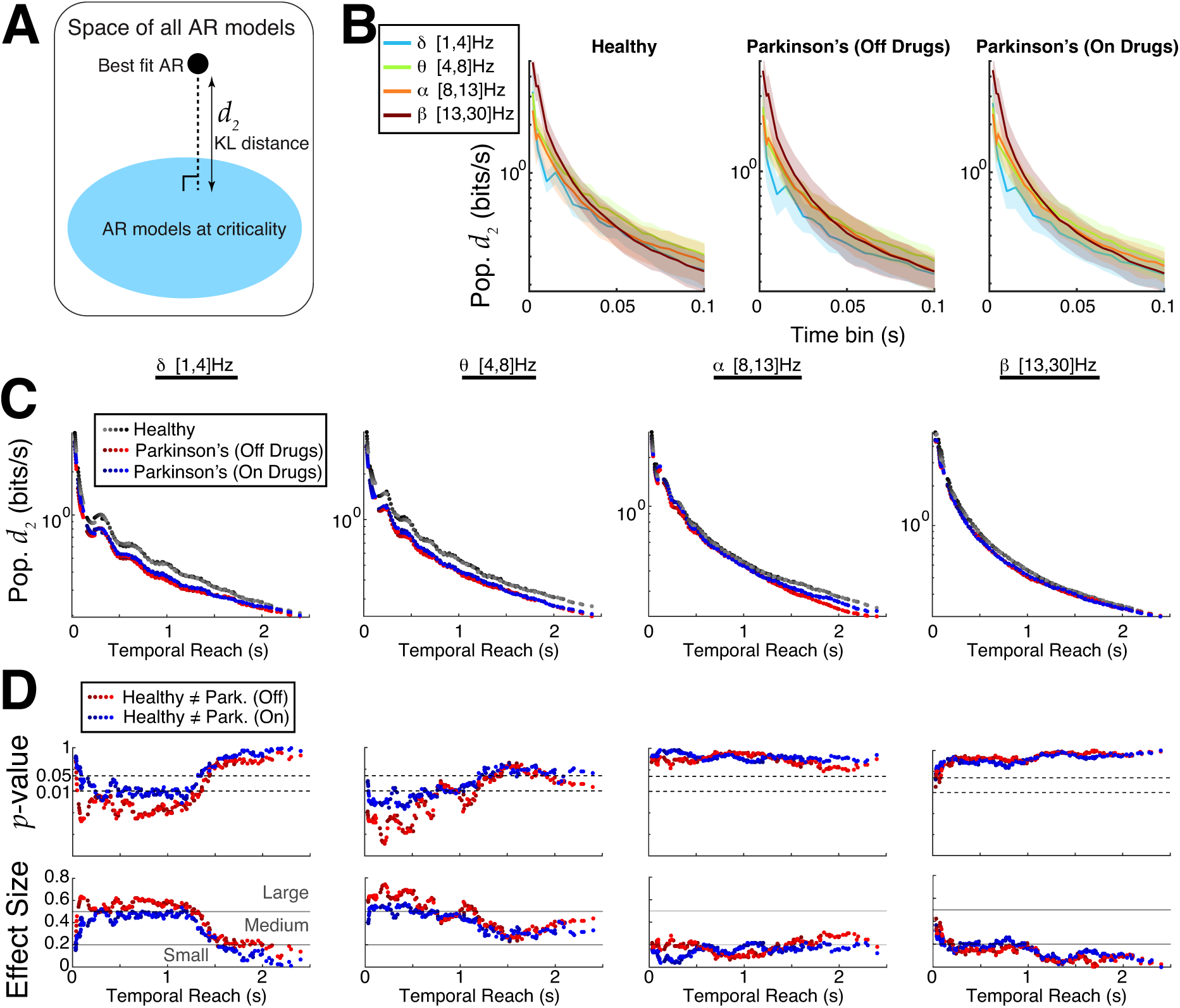
Emergent Parkinson’s oscillations are closer to criticality. **A)** Using new theory to quantify differences in distance to criticality between controls and Parkinson’s patients after data is fit with an AR model. **B)** The population *d*_2_ (bits/s) values (log-scale) grouped by control and two Parkinson’s state as a function of coarse-grained time bin with AR model order 20 shows little difference across different band-passed frequencies. **C)** Summary of population *d*_2_ values (log-scale) for many time bins and model orders; the x-axis represents the ‘temporal reach’, i.e., model order multiplied by time bin length (varies from 2 ms to 100 ms). The control subjects consistently had larger *d*_2_ and were thus further from criticality than Parkinson’s patients, independent of model order, time bin length, or frequency band. **D**) Quantifying the statistical significance of our results using Wilcoxon rank-sum test, showing the *p−*values (log-scale) and effect sizes (see **Methods: Wilcoxon Rank-Sum Test**). The different shades of colors in C) and D) correspond to AR model fits of different orders ranging from 16 to 24.

**Table 1.** Statistics to show that control subjects have shorter time scales than Parkinson’s using ACF timescale measure; see Figure 3C. Using Wilcoxon rank-sum test where the null hypothesis is that both data samples are drawn from the same distribution. Top shows *p−*values, bottom shows effect size (see **Methods: Wilcoxon Rank-Sum Test**) with a qualitative assessment of small, medium, large (Cohen, 2013; Tomczak and Tomczak, 2014).

| Relationship / $p$-value | $\delta$ band | $\theta$ band | $\alpha$ band | $\beta$ band |
| --- | --- | --- | --- | --- |
| Cntrl. vs. Park. (Off drugs) | $1.5 \times 10^{-2}$ | $4.6 \times 10^{-5}$ | 0.33 | 0.18 |
| Cntrl. vs. Park. (On drugs) | $6.6 \times 10^{-2}$ | $2.2 \times 10^{-3}$ | 0.42 | 0.24 |
| (Park.) On vs. Off | 0.43 | 0.27 | 0.91 | 0.88 |
| Relationship / Effect Size | $\delta$ band | $\theta$ band | $\alpha$ band | $\beta$ band |
| Cntrl. vs. Park. (Off drugs) | 0.44 (med) | 0.73 (lrg) | 0.41 (n/a) | 0.24 (n/a) |
| Cntrl. vs. Park. (On drugs) | 0.33 (med) | 0.55 (lrg) | 0.15 (n/a) | 0.21 (n/a) |
| (Park.) On vs. Off | 0.14 (n/a) | 0.13 (n/a) | $2.3 \times 10^{-2}$ (n/a) | $2.7 \times 10^{-2}$ (n/a) |

**Table 2.** Statistics to show that control subjects have shorter time scales than Parkinson’s using DFA coefficient; see Figure 4B. Using Wilcoxon rank-sum test where the null hypothesis is that both data samples are drawn from the same distribution. Top shows *p−*values, bottom shows effect size (see **Methods: Wilcoxon Rank-Sum Test**).

| Relationship / $p$-value | $\delta$ band | $\theta$ band | $\alpha$ band | $\beta$ band |
| --- | --- | --- | --- | --- |
| Cntrl. vs. Park. (Off drugs) | $9.4 \times 10^{-3}$ | $9.4 \times 10^{-3}$ | $8.6 \times 10^{-2}$ | $2.8 \times 10^{-2}$ |
| Cntrl. vs. Park. (On drugs) | $7.7 \times 10^{-2}$ | $5.7 \times 10^{-3}$ | $2.4 \times 10^{-2}$ | $9.1 \times 10^{-3}$ |
| (Park.) On vs. Off | 0.28 | 0.51 | 0.89 | 0.96 |
| Relationship / Effect Size | $\delta$ band | $\theta$ band | $\alpha$ band | $\beta$ band |
| Cntrl. vs. Park. (Off drugs) | 0.47 (med) | 0.47 (med) | 0.31 (med) | 0.11 (sm) |
| Cntrl. vs. Park. (On drugs) | 0.32 (med) | 0.51 (lrg) | 0.42 (med) | 0.14 (sm) |
| (Park.) On vs. Off | 0.2 (n/a) | 0.13 (n/a) | 0.26 (n/a) | $1.5 \times 10^{-3}$ (n/a) |

**Table 3.** The side of reported physical impairment in Parkinson’s patients (Appelhoff et al., 2019; Rockhill et al., 2021) and thus corresponding electrode used. Electrode C3 is on the left motor cortex, C4 on the right motor cortex. Note that all subjects had the same side for physical impairment on and off drug treatment except for subject 14 who switched to Right side (C3) while on drug treatment.

| Subject | Impairment Side | Electrode |
| --- | --- | --- |
| 1 | Right | C3 |
| 2 | Both | Both |
| 3 | Right | C3 |
| 4 | Right | C3 |
| 5 | Left | C4 |
| 6 | Right | C3 |
| 7 | Left | C4 |
| 8 | Left | C4 |
| 9 | Left | C4 |
| 10 | Right | C3 |
| 11 | Right | C3 |
| 12 | Right | C3 |
| 13 | Left | C4 |
| 14 | Left* | C4* |
| 15 | Right | C3 |

Next we perform DFA analysis, which characterizes how fluctuations vary across different timescales, also known as LRTC. In the DFA analysis we find that control subjects have on average shorter range temporal correlation than Parkinson’s patients, consistent with the lower frequency bands in the ACF timescale analysis. A demonstration of the DFA method is depicted in Figure 4A on a control subject’s resting state EEG in the delta-band where the fluctuation amplitude as a function of time window length in log-log coordinates requires 2 lines at a dividing point; such a dividing point is required in about 72% to 85% of the time (counting all frequency bands, subject type, and electrode combinations). In such cases, we use the slope of the best fit line for larger time windows (second segment to the right), and call this the DFA coefficient. When there are no temporal correlations (i.e., white noise) the DFA coefficient is 0.5. In a random walk, temporal memory is infinite and the DFA coefficient is 1.5. DFA coefficients between 0.5 and 1.5 indicate intermediate temporal correlations. A summary of all DFA coefficients is shown with box plots in Fig 4B with four frequency bands: on average control subjects have a much shorter range of temporal correlation, i.e., timescales, than Parkinson’s patients (on or off drugs). The trend that control subjects have shorter timescales than Parkinson’s patients is robust across all four frequency bands we consider, with the alpha-band results having comparatively weaker results with larger *p−*values using Wilcoxon rank-sum test. The DFA coefficients shown include all 16 control subjects, but for Parkinson’s 1 or 2 patients were excluded (depending on frequency band) because the fluctuation amplitudes for a few subjects were too variable to be well fit by a line (see Fig S3 and Supplementary Text S1). Figure 4C clearly demonstrates how different the population averages are; control subjects have much smaller average DFA coefficients than Parkinson’s, and Parkinson’s patients have similar DFA coefficients regardless of whether on or off drug treatments.

Although both ACF and DFA results provide evidence that control subjects’ EEG motor cortex activity is further from criticality than Parkinson’s patients, these analyses do not directly measure distance to criticality. To this end, we use our recently developed rigorous theory to directly calculate distance to criticality (*d*_2_). The *d*_2_ measure has several specific advantages : i) unlike DFA, it does not require specifically choosing a time window segment and assessing quality of linear fits (Fig S3), ii) unlike ACF, it does not require a prescribed threshold to find timescale, iii) the distance to criticality *d*_2_ (Fig 5A) is a precise quantification of distance independent of model parameterization, calculated in units of bits/sec (measure of evidence for ruling out being at criticality). Our framework requires first fitting an auto-regressive (AR) model to the data, then calculating the distance of the fitted model to the critical state; see **Methods: Temporal Renormalization Group Theory** and Figs S5–S7 for further details.

We perform a detailed comparison of Parkinson’s and control subjects using *d*_2_ and find control subjects’ EEG in primary motor cortex are generally further from criticality than Parkinson’s patients. We specifically vary the AR model order (16 to 24) as well as the time bin length (2 ms up to 100 ms) – we previously showed that increasing the time bin can unveil critical dynamics (Fontenele et al., 2024) and that *d*_2_ is expected to decrease monotonically with increasing model order and with increasing time bin length (Sooter et al., 2026). Figure 5B shows, within a given subject group (healthy, Parkinson’s on/off drugs) for a fixed AR model order 20, that *d*_2_ tends to decrease as time bin length increases, except occasionally in the delta-band (light blue), and that there are minor differences in population *d*_2_ across frequency bands with a given time bin. Varying AR model order and time bin length simultaneously results in a wide variety of ‘**temporal reach**’ (x-axis of Fig 5C,D) that we define as the the AR model order multiplied by the specific time bin, i.e., the maximal time in the past that can influence the present AR model value. The temporal reach values we consider have a large range from 32 ms to 2.4 s. The population averaged *d*_2_ for many temporal reach values is shown in Figure 5C (y-axis is log-scale, different color shades correspond to different AR model order), where it is evident that Parkinson’s patients (red and blue dots) are closer to criticality (*d*_2_ below) than controls (black/gray) in the delta- and theta-bands. We use Wilcoxon rank-sum test to analyze whether differences between Parkinson’s and control are statistically significant (*p−*values in top row of Fig 5D). The differences are clear in the delta- and theta-bands for a wide range of temporal reach values, there is no differences in the alpha band, and differences in the beta-band are only evident with small temporal reach values. The effect size and a qualitative characterization of effect size is shown in the bottom row of Figure 5D.

## Discussion

Here we have shown that the prominent delta- and theta-band oscillations that emerge in Parkinson’s disease from resting state EEG motor cortex are, in fact, near-critical oscillations. Although each of these oscillations is defined by particular timescales (the oscillation periods), the power (amplitude) of these oscillations exhibits fluctuations across a wide range of time scales. These amplitude fluctuations are approximately scale invariant, which is how critical oscillations are defined (Fontenele et al., 2026; Palva and Palva, 2018). In contrast, in healthy controls, the same frequency bands have amplitude fluctuations that are further from criticality.

The distance measure *d*_2_ enables a fair comparison of different time series, and is a rigorous information-theoretic entity in units of bits/s that measures the amount of evidence for ruling out the hypothesis that the data are at criticality (Sooter et al., 2026). Although the ACF and DFA analyses yielded similar results, *d*_2_ is a direct measure for distance to temporal scale-invariance, and proved to be cleaner for delineating differences (control vs. Parkinson’s), and did not require making specific choices regarding threshold cut-offs, which time window segments to use, etc. Unlike traditional methods, the analysis with *d*_2_ goes beyond just measuring timescale and has implications on neural computation (Hengen and Shew, 2025). For both DFA and *d*_2_, the strongest separation between control and Parkinson’s patients are in the delta- and theta-bands, followed by the beta-band, with the weakest results in the alpha-band where oscillations are known to be a sign of healthy function (Britton et al., 2016). The ACF timescales only had significant differences in the delta- and theta-band. The relative consistency of results for these different methods suggests that there are real and surprising differences between control and Parkinson’s patients in motor cortex EEG. In awake mice models of Parkinson’s disease, recent studies have shown the emergence of delta oscillations in parts of the basal ganglia (substantia nigra spiking) that are connected to motor cortex (Aristieta et al., 2024; Whalen et al., 2020); with an accompanying computational study to show that delta oscillations can be transmitted to different regions in the basal ganglia (Whalen et al., 2023). All together these studies suggest the emergence of lower frequency band oscillations in Parkinson’s patients should not be overlook as a crucial signature of disease despite the ubiquity of emergent beta-band oscillations (Brown et al., 2001; Sharott et al., 2005; George et al., 2013).

Our results in motor cortex are at odds with the idea that, in general, healthy brains operate closer to criticality than pathological ones (Hengen and Shew, 2025; O’Byrne and Jerbi, 2022), as discussed in the Introduction. However, our results are in line with two recent publications where it was reported that Parkinson’s patients i) have more frequency bands closer to ‘edge of chaos’ (a point related to criticality) than control subjects in whole brain MEG (Calvo et al., 2024), and ii) can have longer timescales as measured with DFA coefficients in whole brain EEG in the theta-band (Lee et al., 2024). These studies are different than ours because they focused on whole brain imaging and included many more subjects with which they aggregated/averaged. The number of subjects we used enabled detailed analysis, for example to assess in detail the quality of model fits for each subject.

Along these lines, another recent study showed that a measure of ‘intrinsic neural timescale’ using fMRI was longer in late stage Parkinson’s patients than in healthy controls in the anterior cortical region (Wei et al., 2024). This study is in line with our results, but unlike our *d*_2_, their measure of intrinsic neural timescale is indirect because it involves calculating when various ACFs first cross below a threshold, smoothing the maximum of those values over space and applying a z-transform. The timescales of fMRI measurements are coarser than those of EEG, with temporal resolution on the order of seconds, so we cannot make any direct comparisons with our results. Moreover, our motor cortex EEG results focuses on only one piece of the circuit. More detailed studies to connect spiking neural network mechanisms to critical dynamics in EEG have been addressed (Brake et al., 2024) but, to our knowledge, not directly in Parkinson’s disease.

Interestingly, Parkinson patients on versus off drugs to treat motor symptoms did not ever have statistically significant differences in their motor cortex EEG, independent of the methods (ACF, DFA, *d*_2_). Presumably, these drugs helped mitigate their motor symptoms to some extent, but the motor cortex activity that is responsible for voluntary movement planning and muscle control did not exhibit any changes in timescales <u>in the resting state</u>. Perhaps EEG amplitude fluctuations in motor cortex in the resting state are not a direct reflection of mitigated motor symptoms with medication, but rather indicate the difference between Parkinson’s disease itself compared to control.

## Methods

### Ethics statement

This article presents an accurate account of the work performed by the stated authors, and all underlying data are represented accurately with consent from the owners. To the best of our knowledge, this work is original and is not under consideration for publication elsewhere. The study used publicly available data with accurate citation, and all methods were performed in accordance with relevant guidelines and regulations. The authors declare no conflicts of interest related to this research.

This study uses third party human EEG data that is publicly available (George et al., 2013; Swann et al., 2015; Jackson et al., 2019) (see Data and code availability section). The Materials and methods section in their papers explicitly state that ‘All the participants provided written informed consent according to an Institutional Review Board Protocol at the University of California, San Diego and the Declaration of Helsinki’. We have also obtained written approval from the authors to use their data in this study.

### Parkinson’s patient characteristics

Table 3 shows side of physical impairment in the Parkinson’s patients.

### Autocorrelation function

The autocorrelation function is a common tool used to characterize how related (correlated) a time series of data *x*(*t*) is with specific time lags *τ*. The autocovariance function of a time series *x*(*t*) is:

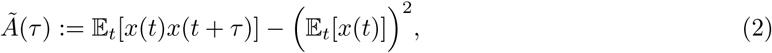

and the autocorrelation function is simply:

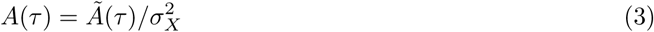

where 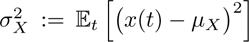 is the point-wise variance of the time series. We calculate the auto-correlation function of a particular time series of EEG via the Matlab function *autocorr* on centered data 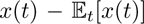 with 10,000 lags (recall the time bins are 2 ms) and 1.96 standard deviations: *autocorr(X-mean(X),‘NumLags’,10000,‘NumStd’,1.96)*. The results are in Figures 3, S1.

### Detrended Fluctuation Analysis (DFA)

DFA is a common method for quantifying the degree of long-range temporal correlations (Peng et al., 1994). For a given time-series, the DFA coefficient was calculated by assessing the correlation of fluctuation amplitudes in various time window lengths. Start with a time-series x*_j_*, then calculate the cumulative sum: 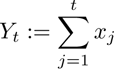. The ‘entire’ *Y_t_* time-series is divided into *n* equal lengths for the duration of the specified time window *τ_x_* of length (*τ_x_/dt* + 1) – if the length of *Y_t_* cannot be evenly divided, the end of the time-series is truncated, so *n* = *⌊N/*(*τ_x_/dt* + 1)*⌋*. Then for each segment of length (*τ_x_/dt* + 1) the *local trend* (least squares linear fit *L_k_*) is calculated. After which the mean-squared deviation is calculated: 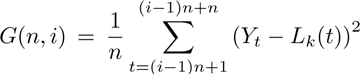, then the mean fluctuation amplitude is: 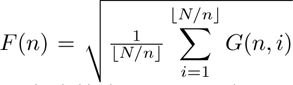. Finally, the least squares linear fit between log(n) (horizontal axis) and log(*F* (*n*)) (vertical axis) is calculated – the slope of this line is called the *DFA coefficient*.

In cases where log(n) versus log(*F* (*n*)) is not well-fit by a single line, the time windows are split in two segments determined manually, then two least squares linear fits are calculated with the larger windows (right half) determining the DFA coefficient (Gu et al., 2015).

### Temporal renormalization group (tRG) theory

A system is at criticality if (1) it lies at a boundary between qualitatively different operating regimes and (2) it exhibits scale-invariance, i.e. the lack of a characteristic spatial or temporal scale (Hengen and Shew, 2025). The renormalization group (**RG**), which was originally developed to study critical phenomena in condensed matter systems, brings mathematical precision to these statements. The core idea of RG is to gradually remove the fine-scale details of a model to generate new, effective models at coarser scales. Fixed points of the RG operation therefore correspond to models that are scale-invariant, and all of the models in the basin of attraction of such a fixed point share the same coarse-scale behavior - this is the fundamental reason why, for example, water and ferromagnets poised near their respective phase transitions have quantitatively identical scaling exponents despite their drastic dissimilarities at microscopic length scales. Some RG fixed points are stable, meaning that there is an extended region in model space surrounding the fixed point such that every model in the region flows into the fixed point. Such fixed points fail to satisfy condition (1) in the definition of criticality. (For example, the RG fixed point corresponding to the disordered phase of an Ising model is stable.) Models lying in the basins of attraction of *unstable* fixed points, on the other hand, are both scale-invariant and poised at a boundary between different operating regimes, and hence are at criticality.

In traditional applications of RG to spatially organized systems (e.g. Ising-type models), coarse-graining is implemented in space. Neural systems, on the other hand, can have rich temporal dynamics in an measurable entity (i.e., population EEG) independent of whether there is weak or crucial spatial structure. In Sooter et al. (2026), we argued that the appropriate way to define criticality in such systems is with a temporal RG (**tRG**), wherein high-frequency features of a model are gradually removed to reveal its asymptotic behavior at low frequencies. We applied this procedure to a fundamental class of univariate discrete-time stochastic dynamical systems, Gaussian autoregressive (**AR**) models:

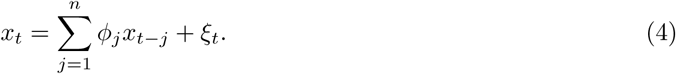

We chose AR models both because they are analytically tractable, and because they are the optimal choice in a precise *max ent* sense. Specifically, if an observed time series is short enough that we can only confidently estimate its second-order (autocovariance) statistics, then AR models are the maximum entropy (i.e. minimally presumptive) way to model those statistics (Choi and Cover, 1984). In an AR model, the state x*_t_* at time t is a linear, Gaussian readout of the recent history (up to some maximum lag n, called the model order):

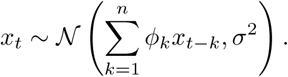

In the space of order-*n* AR models, there are *n* + 1 tRG fixed points, which we can label using the power-law exponents of their respective power spectra, *β* = 0, 2*, …,* 2*n*. The *β* = 0 fixed point is stable and corresponds to white noise - this is the “trivial” fixed point that any AR model with a finite characteristic timescale flows into. The basins of attraction of the *β ≥* 2 fixed points constitute the AR models that are at criticality.

Next, we asked how we should quantify proximity to these basins. That is, having determined which AR models are *at* criticality, can we say which ones are *close* to criticality? Naively, we could measure the Euclidean distance (in the parameter space defined by the AR model history kernel *ϕ*) from an AR model to each of the basins of attraction. However, there is no principled reason to use Euclidean distance rather than some other metric. To resolve this ambiguity, we turned to information theory and *define* proximity to criticality as distinguishability (per unit time) from a system at criticality. Specifically, for a given order-*n* AR model *B*, let:

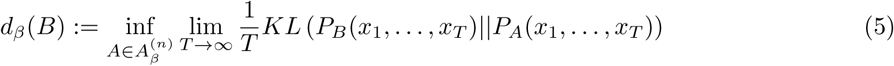

where 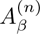 is the set of order-*n* AR models that flow into the *β* fixed point, *P_A_*(*x*_1_*, …, x_T_*) is the probability distribution for a *T −*step draw from the AR model *A* (and similarly for *P_B_*), and *KL*(·||*·*) is the Kullback-Leibler divergence. The structure of basins of attraction is such that that infimum taken over the set of all critical models 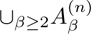 is equal to the infimum taken over the *β* = 2 basin of attraction (Sooter et al., 2026); hence we only report *d*_2_ in this paper.

To estimate *d*_2_ from EEG data after bandpass filtering and extracting the envelope, we: 1) fit an AR model to the data using the Yule-Walker method, and 2) compute *d*_2_ for this model using Eq (5).

### Wilcoxon Rank-Sum Test

We use the Wilcoxon rank-sum test (**WRST**) because it is ideal for the EEG. It is a nonparametric test of the null hypothesis that two groups of data are generated from the same distribution. The *p−*values of this test correspond to the probability that the null hypothesis holds. In addition, we report the *Effect Size* of the WRST:

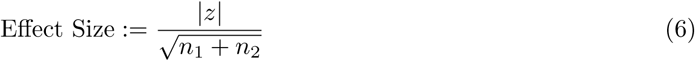

where *z* is the *z−*score of the *U−*statistic, *z* = (*U − µ_U_*)*/σ_U_* and *n_j_* are the sample sizes for the two populations. Effect sizes for *t−*test and Wilcoxon rank-sum test with values: (0, 0.2] are considered **small**, (0.2, 0.5] are **medium**, (0.5, 0.8] and above are **large** (Cohen, 2013; Tomczak and Tomczak, 2014); these labels are simply a qualitative assessment.

### EEG Data

We used freely available EEG collected years ago that have appeared in many studies (Jackson et al., 2019; Swann et al., 2015; George et al., 2013) and was made widely applicable following common standards (Pernet et al., 2019; Appelhoff et al., 2019). We used all 16 control (control) subjects and all 15 Parkinson’s patients except for some of the DFA coefficients (see Figure S3). We used an EEG reader function by Tcheslavski (2025).

### Frequency Band Limits

The frequency bands of interest were limited with an upper bound in the beta band (30 Hz) because the 60 Hz grounding frequency (inferred by the power spectrum of all electrodes have a large peak at 60 Hz); any signals close to 60 Hz are considered artifacts perhaps due to electrical interference. For completeness and since some EEG studies report results in the gamma band frequency (George et al., 2013; Swann et al., 2015), we repeated our analysis on a lower gamma band frequency between 30 and 50 Hz (as was done in George et al. (2013), and found that the general trend of results we observed did not hold (see Fig S4, except for the population averaged ACF decay Fig S4A). We note however that in this lower gamma frequency band, the DFA analysis was messier than the other 4 lower frequency bands; in particular for the control subjects where 5 were excluded (see GitHub page), and in one of the ACF timescales was unusually long (longer than 20 s). Thus, these results should be taken with caution.

## Data and code availability Declarations

### Data availability

The raw EEG dataset was collected at UC San Diego from a team of researchers, it is freely available (Rockhill et al., 2021) at https://openneuro.org/datasets/ds002778/versions/1.0.2.

### Code availability

See https://github.com/chengly70/parkeeg for MATLAB code implementing all computational components in this paper.

### Author Contributions

Conceptualization: CL, WLS. Methodology: CL, JSS, WLS. Software: JSS, AJF, CL. Validation: CL. Formal Analysis: AJF, CL. Investigation: JSS, CL. Resources: N/A. Data Curation: CL. Writing original draft: CL, WLS, JSS, AKB. Writing review and editing: CL, JSS, AKB, AJF, WLS. Visualization: AJF, CL. Supervision: CL. Project administration: CL. Funding acquisition: CL, JSS, AKB, WLS.

