## Supplementary Material for "Emergent critical oscillations in motor cortex of Parkinson’s patients"

### Similar results when using both electrodes

This section shows the analogous results in the main text except for the Parkinson’s patients, we use the average of both C3 and C4 electrodes without regard to the side of physical impairment shown in Table 3.

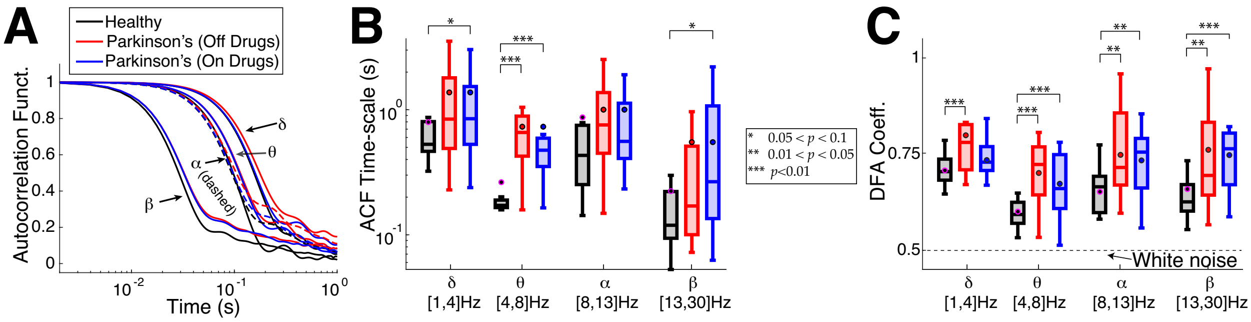


Figure S1: ACF and DFA results using both C3 and C4.
**A**) The population averaged ACF. **B**) The ACF has the same trend as before, with control subjects decaying faster than Parkinson’s for many frequency bands. The dots on the box plot with black (or magenta) outline represent the population mean here and in panel C. **C**) The control subjects generally have smaller DFA coefficients and are further from criticality than Parkinson’s patients, however the results here in the $\delta$ and $\theta$ band are not as strong as in the main text (the $p-$values for the differences are larger). But the results are stronger in the $\alpha$ band compared to the main text (small $p-$values, see Table S1). **C**) Differences in the population average DFA coefficients. See Figures 2–3 for comparison.

Table S1: **Statistics to show that control subjects have shorter time scales than Parkinson’, using DFA coefficient of average of both electrodes; see Figure S1B**. Using Wilcoxon rank-sum test where the null hypothesis is that both data samples are drawn from the same distribution. Top shows $p-$values, bottom shows effect size (see **Methods: Wilcoxon Rank-Sum Test**).

| Relationship / $p-$value | $\delta$ band | $\theta$ band | $\alpha$ band | $\beta$ band |
|---|---|---|---|---|
| Cntrl. vs. Park. (Off drugs) | $8.3 \times 10^{-3}$ | $1.3 \times 10^{-3}$ | $2.2 \times 10^{-2}$ | $4.0 \times 10^{-2}$ |
| Cntrl. vs. Park. (On drugs) | $1.3 \times 10^{-1}$ | $3.2 \times 10^{-3}$ | $1.7 \times 10^{-2}$ | $9.1 \times 10^{-3}$ |
| (Park.) On vs. Off | 0.18 | 0.27 | 0.90 | 0.98 |

| Relationship / Effect Size | $\delta$ band | $\theta$ band | $\alpha$ band | $\beta$ band |
|---|---|---|---|---|
| Cntrl. vs. Park. (Off drugs) | 0.48 (med) | 0.60 (lrg) | 0.43 (med) | 0.38 (med) |
| Cntrl. vs. Park. (On drugs) | 0.28 (med) | 0.56 (lrg) | 0.45 (med) | 0.48 (med) |
| (Park.) On vs. Off | 0.26 (n/a) | 0.51 (n/a) | 0.023 (n/a) | $4.7 \times 10^{-3}$ (n/a) |

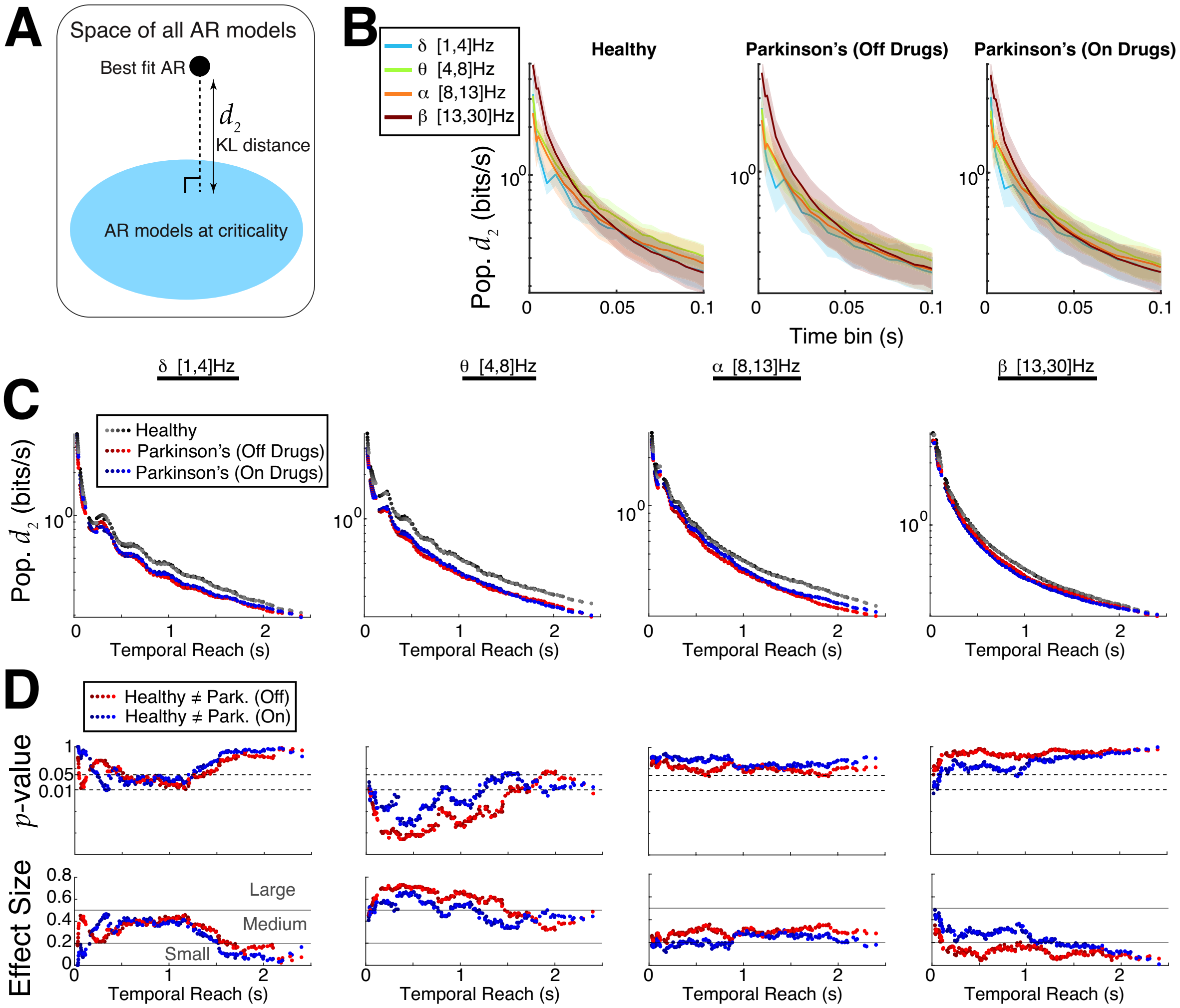


Figure S2: Distance to criticality $d_2$ results using both C3 and C4.
The results here are so similar to before (Fig 4) that the differences in almost all plots are only discernible with side-by-side comparisons. **A**) Methodology (same). **B**) Within a state (control, Parkinson') the population averaged $d_2$ in different band passes do not vary much (for AR model order 20). **C**) The population averaged $d_2$ shows that control subjects are consistently above Parkinson's patients with the strongest results in the $\delta$ and $\theta$ bands. **D**) The $p-$values and effect sizes using Wilcoxon Rank-Sum Test to rule out the null hypothesis that the distances to criticality $d_2$ are from the same distribution. Compared to Figure 4, the results are not as strong here in the $\delta$ band but are stronger in the $\theta$ band; the $\alpha$ and $\beta$ bands are similar to (or perhaps only very slightly better than) Figure 4.

### Details: linear fits for DFA coefficient, low gamma band results

All control subjects were included in the DFA analysis. But due to insufficient linear trends, we excluded some Parkinson's patients: in delta band in Parkinson-OFF, exclude subject 4, in Parkinson-ON, exclude subject 15; in theta band in Parkinson-OFF, exclude subject 4, in Parkinson-ON, exclude subjects 9 and 15; in alpha and beta band in Parkinson-ON, exclude subjects 9 and 15. See `https://github.com/chengly70/parkeeg`.

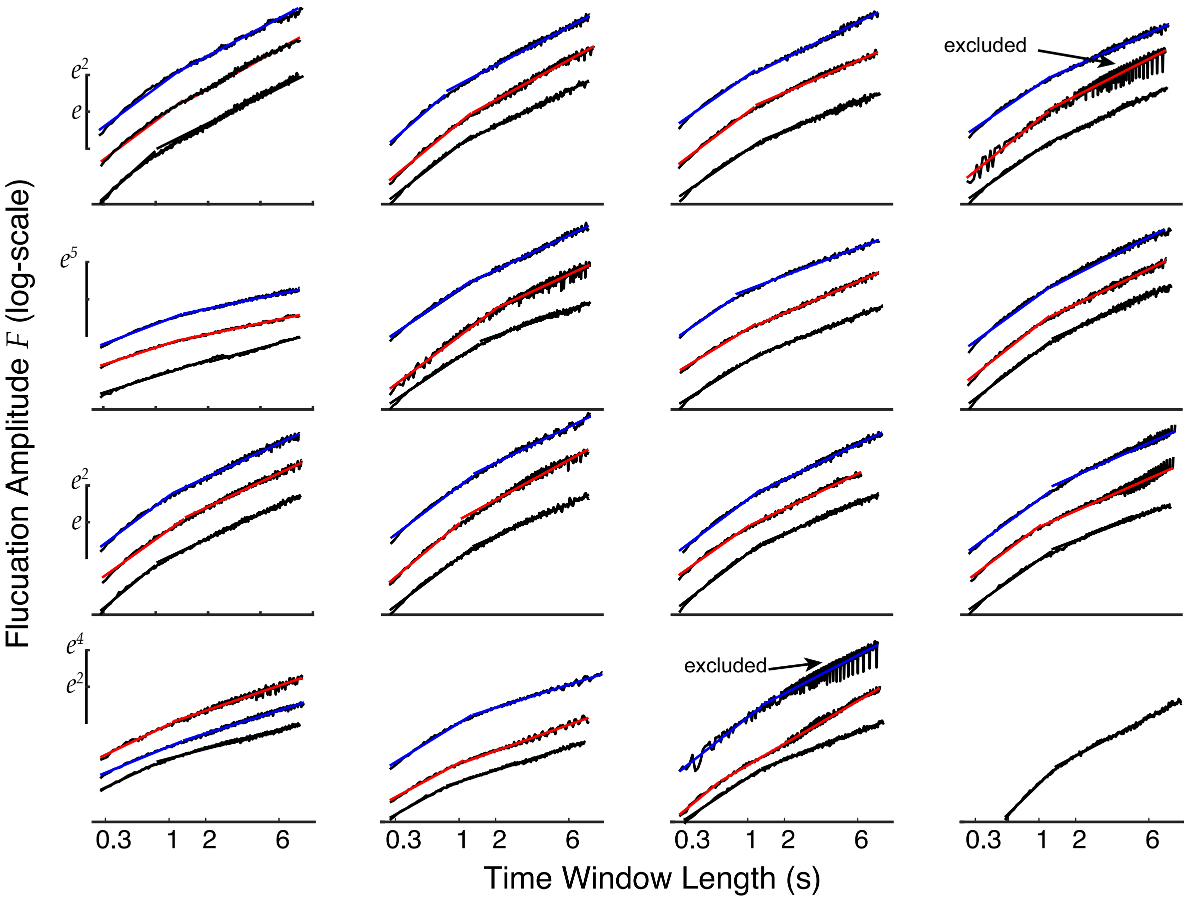


Figure S3: Showing DFA linear fits to all subjects.
Showing fits to the envelope extracted EEG data in $\delta$ band; the other band-passed filtered data fits are similar. The 16 total control subjects are ordered left to right starting at the top row; the 15 Parkinson's patients are plotted (red and blue correspond to off and on drugs, respectively) with the controls for exposition – there is no relationship between the 3 plots on a given set of axes.

### Detailed AR model fits to EEG data

All AR models shown were order 20 for exposition purposes.

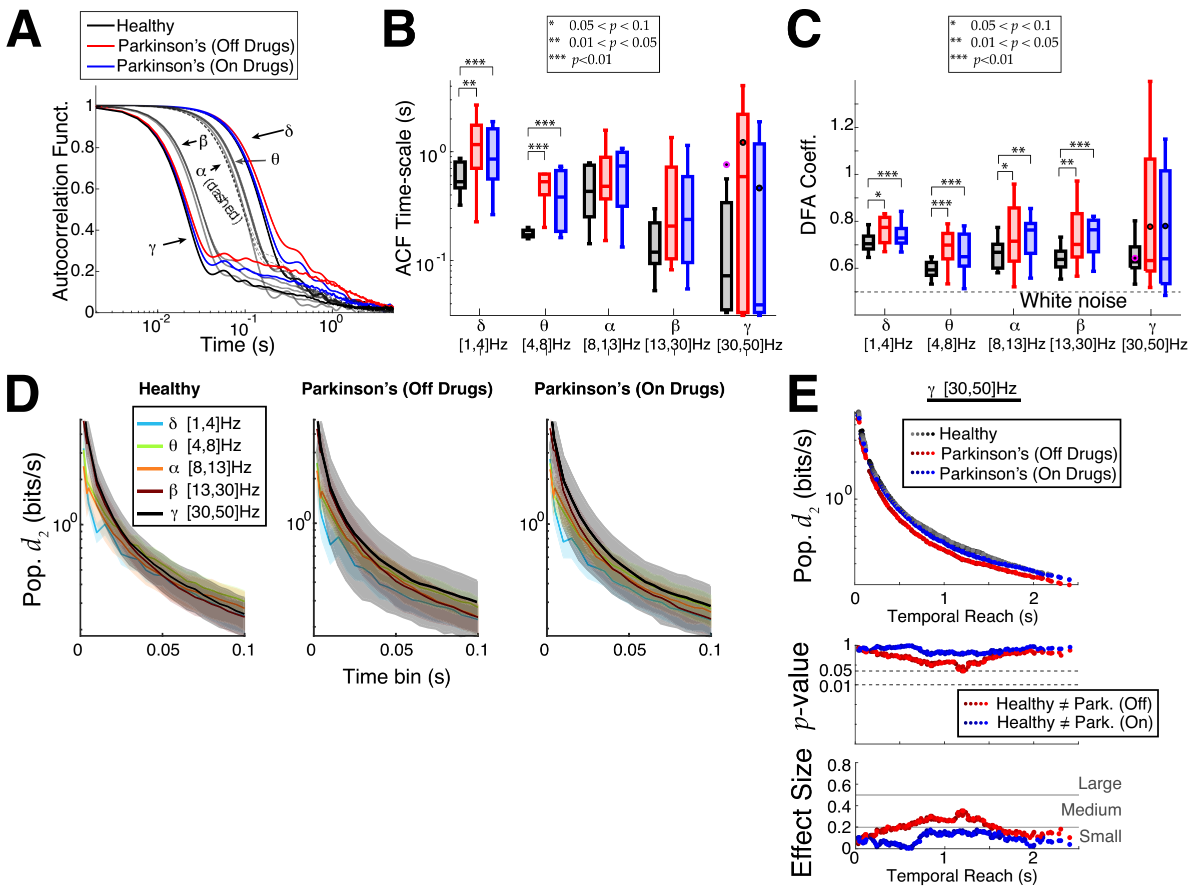


Figure S4: Analyses for low gamma band frequency.

**A**) The ACF trend in Figure 2A holds for the gamma band. **B**) The ACF timescale in the gamma band (far right) is not like the lower 2 frequency bands, there is no statistically significant differences between control and Parkinson's. The dots with black (or magenta) borders correspond to the mean ACF timescale. Subject 14 in the low $\gamma$ band has ACF with slow decay; we set ACF timescale to 11.74 s. **C**) The DFA coefficient in the gamma band (far right) is not like the other bands, there is no statistically significant differences between control and Parkinson's. The Parkinson's patients have higher mean (dots) because of large outlier values but the difference in means are also not statistically significant. **D**) Population averaged $d_2$ for AR model order 20 as the time bin varies shows no distinction within a state (control, etc., see Fig 4B). **E**) In the gamma band, there are no statistically significant differences between control (healthy) and Parkinson's patients, and no statistically significant differences between Parkinson's on versus off drugs.

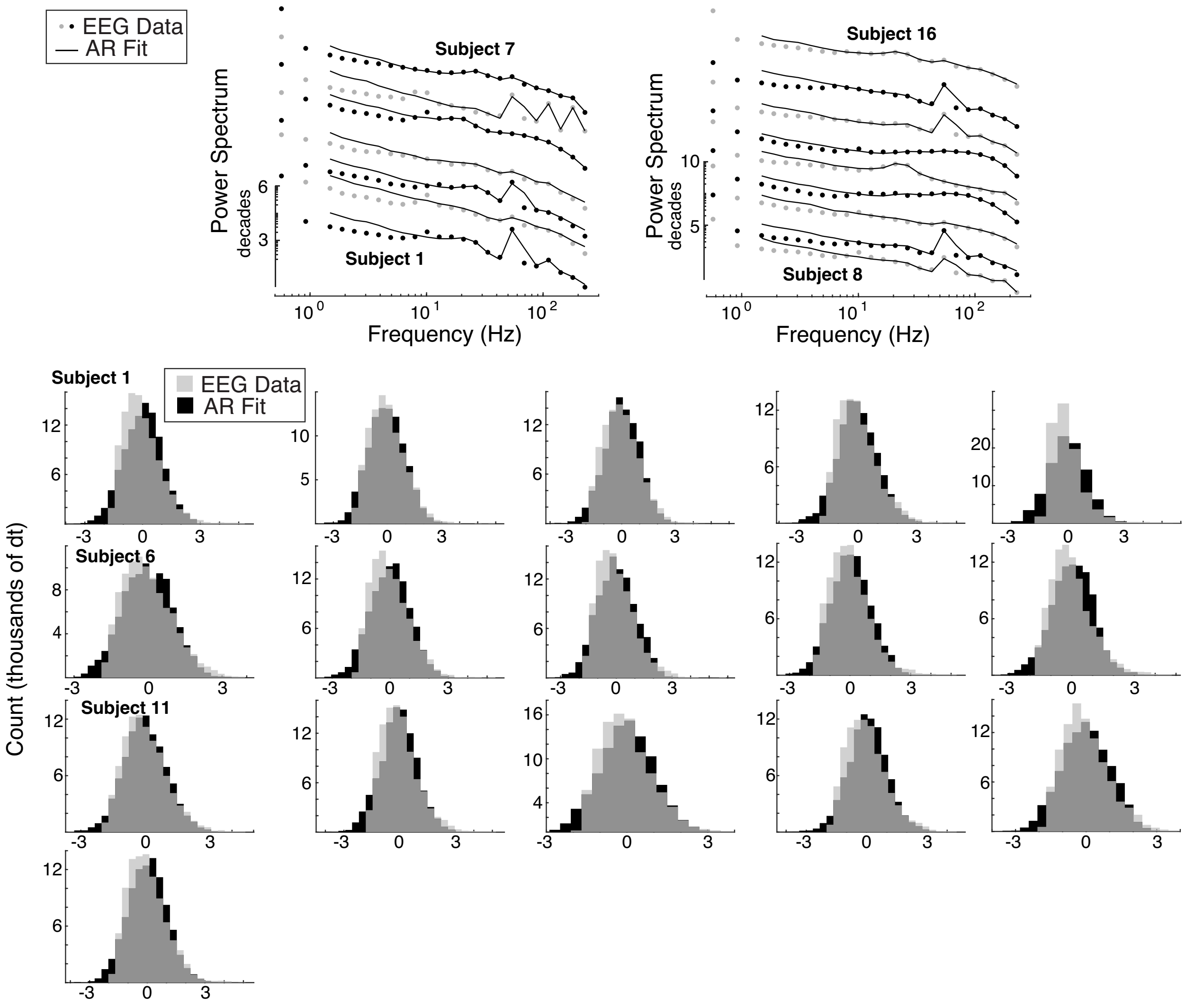


Figure S5: The AR model fits to control subjects' EEG envelope extracted data with finest temporal resolution $dt$. Note that the band-pass filtered data fits with other time bin lengths also perform well. Top half shows the power spectrum for all 16 subjects; EEG with dots, and AR model fit with lines. The bottom half compares the histograms of the $z-$scored point-wise EEG data (gray) and AR model fit (black). Even though the data can be skewed, the (symmetric) AR model fits reasonably well.

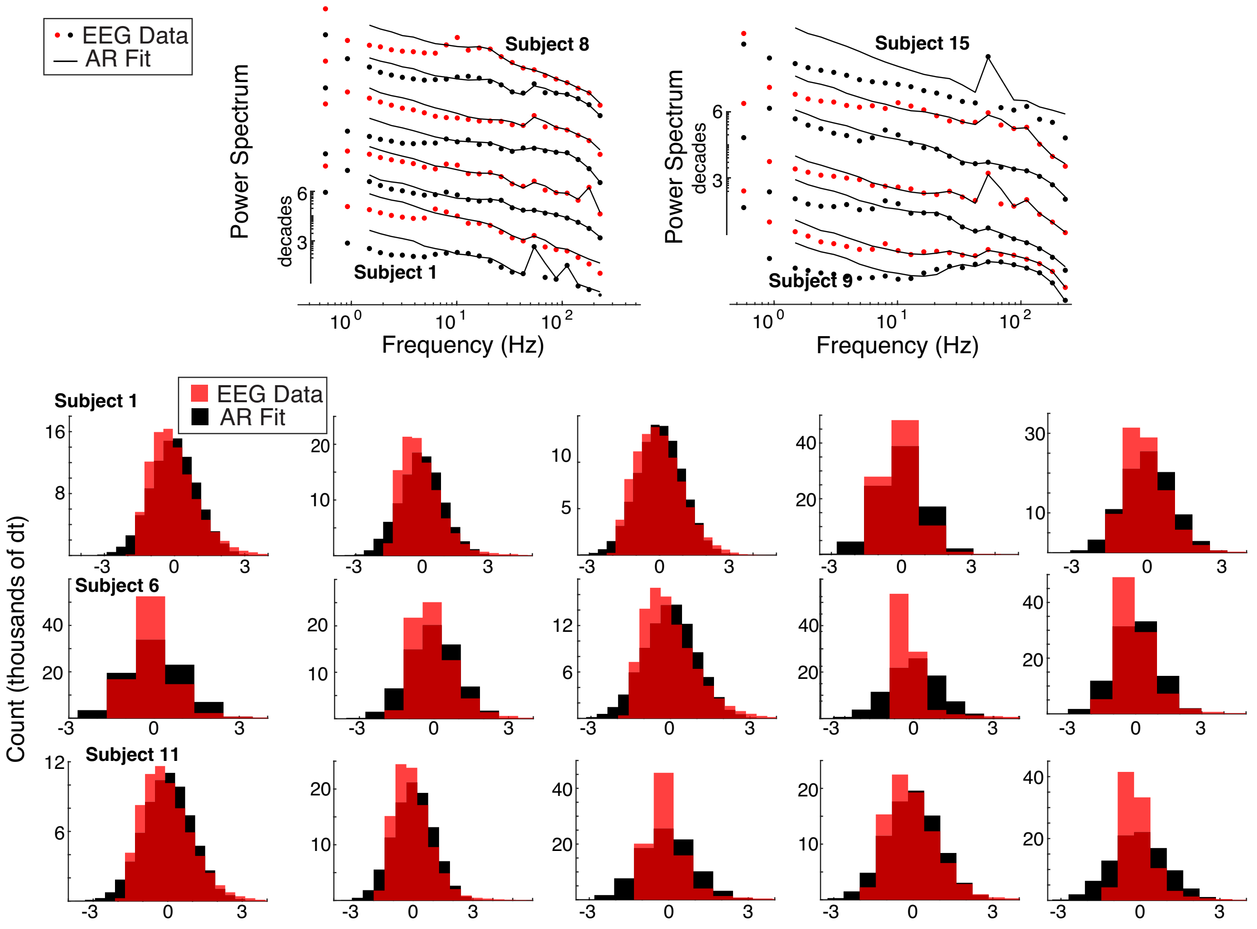


Figure S6: The AR model fits EEG envelope extracted data with finest temporal resolution $dt$ for all 15 Parkinson's patients off medication (same format as Fig S5). The power spectrum (top half) from the data (dots) and AR model fit (lines). The AR model fits (black solid lines) fit the Parkinson's subject data (red and black dots, alternating for visualization) reasonably well. The bottom half compares the histograms of the $z-$scored point-wise EEG data (red) and AR model fit (black).

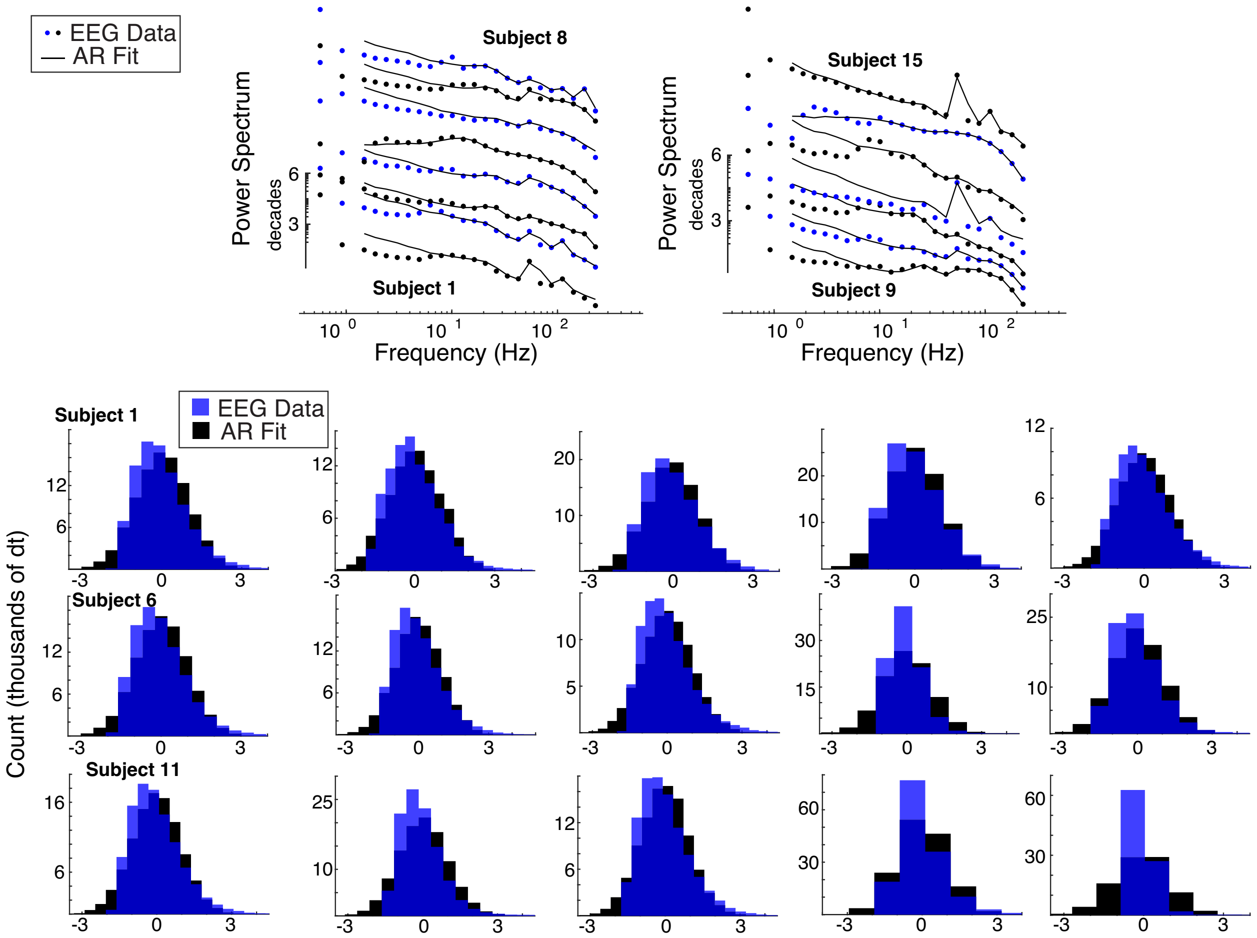


Figure S7: The AR model fits EEG envelope extracted data with finest temporal resolution $dt$ for Parkinson's patients on medication (same format as Figs S5 and S6). Top half show the EEG power spectrum by subject compared (dots) to the AR model fit (line). The bottom half compares the histograms of the $z-$scored point-wise EEG data (blue) and AR model fit (black).
